## Supplementary material for "Temporal selectivity declines in the aging human auditory cortex": Collection of supplementary figures

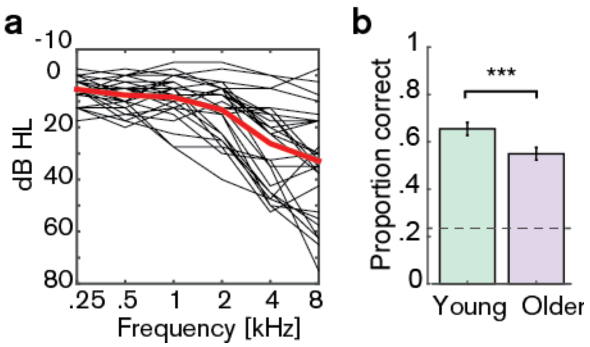


**Figure 1 – figure supplement 1.** (a) **Audiograms** for older participants (averaged over left and right ear). (b) **Average behavioral performance** (mean ± SEM) in the scanner in a four-choice task (3 questions on the story content after each run) for young (green) and older participants (violet). Chance level was 0.25 [proportion correct]. Young participants performed significantly better than older participants (mean proportion correct difference =0.1, $p = 0.009$ [permutation test], Cohen's $d = 0.65$).

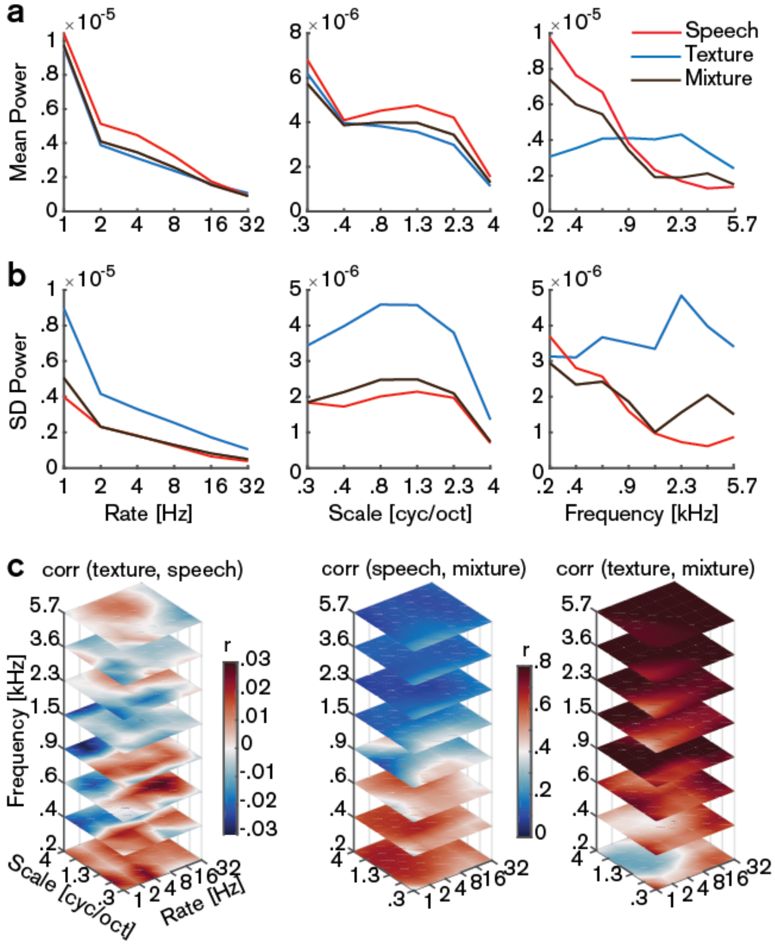


**Figure 1 – figure supplement 2. Modulation spectra of the stimuli.** (a) Average of the marginalized modulation spectra for the stimulus streams separately and their mixture. (b) Standard deviation (SD) of the marginalized modulation spectra. (c) Pairwise correlations (Pearson's *r*) between the different streams in the modulation representation. While speech and texture are mostly uncorrelated over time, the speech–texture mixture is highly correlated with the speech stream in the low frequency range and with textures in the high frequency range. Note that the left plot has a different scaling than the middle and right plot.

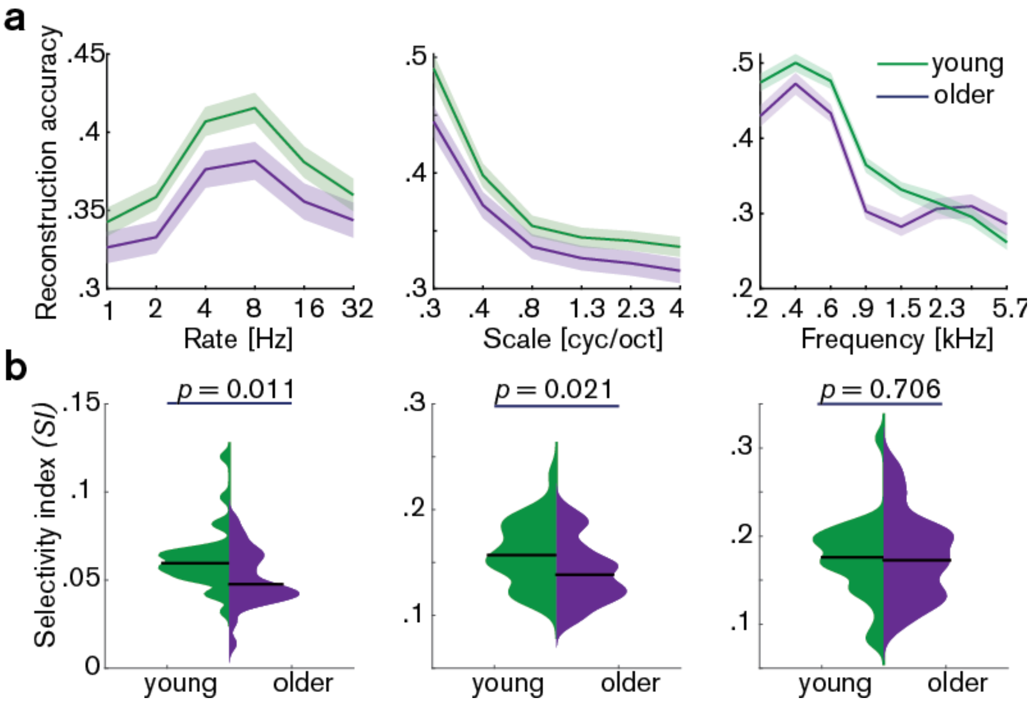


**Figure 5 – figure supplement 1. Decoding from auditory cortex with ICA(AROMA)-cleaned data.** (a) Mean ± SEM of the MTFs' marginal profiles for rate, scale and frequency for young (green) and older (violet) participants separately. (b) The selectivity index *SI* (see Eq. [7]) was compared per acoustic dimension between age groups using an exact permutation test. Note that irrespective of preprocessing (with or without ICA-cleaning, Fig. 5), reconstruction accuracies peak at temporal rates of 4 – 8 Hz. Likewise, for AROMA-cleaned data *SI* for temporal rates is higher in young compared to older participants. Removing the young-subject outlier for rate *SI* (exceeding the grand average by ± 2 SD) does not alter the *SI* mean age difference qualitatively ($p = 0.027$).

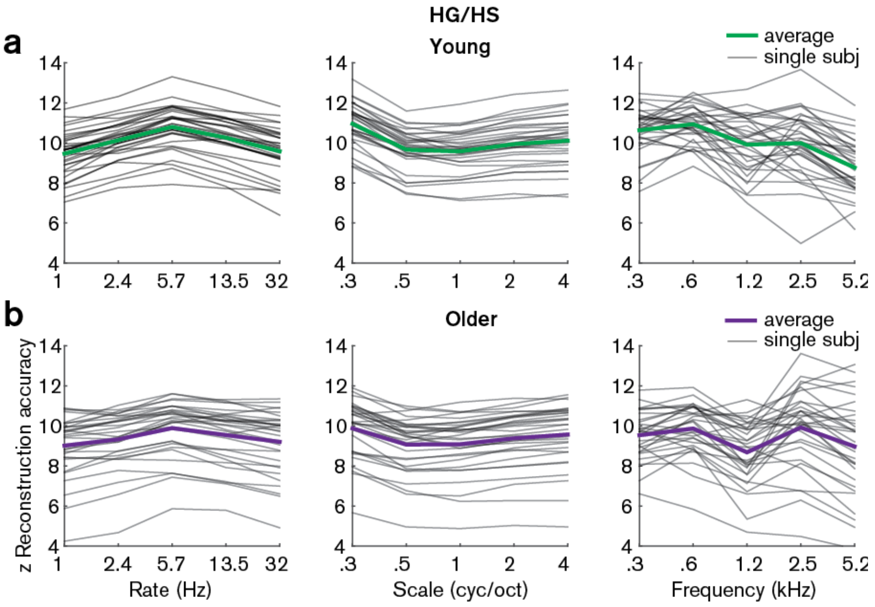


**Figure 7 – figure supplement 1. Z-scored reconstruction accuracies from Heschl's gyrus/sulcus.** Single subject and median reconstruction accuracy profiles for temporal rate, spectral scale and frequency z-scored with respect to the empirical null distribution for young (a) and older (b) participants. Note that all z-values are > 1.96 and are thus significant at $p < 0.05$.

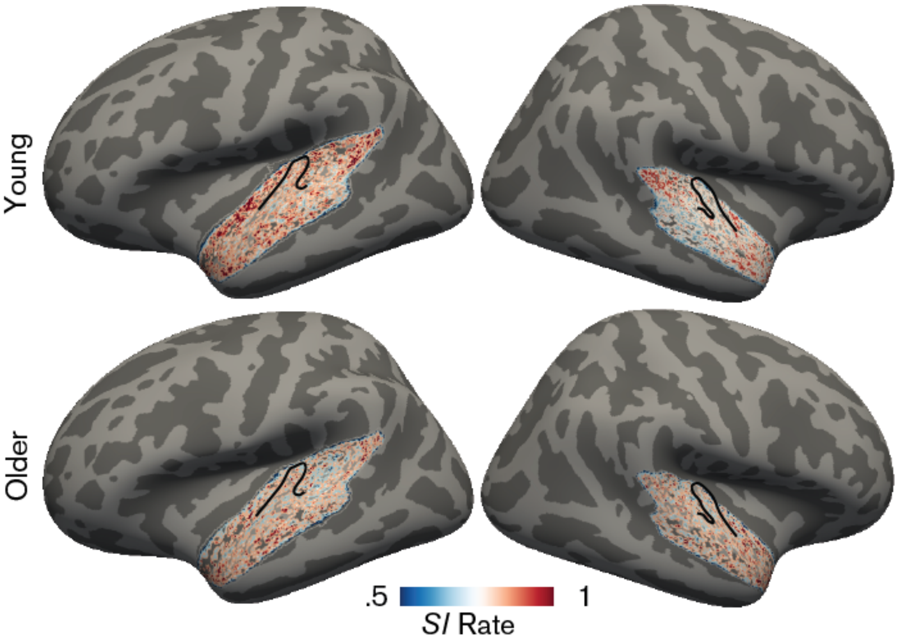


**Figure 7 – figure supplement 2. Topographic maps for the rate selectivity index.** Maps were derived by calculating the *SI* across the rate profile for a given voxel and averaged in the young group (upper panel) and older group (bottom panel). Higher *SI* (red) indicates higher selectivity. Black outlines indicate Heschl's gyrus.

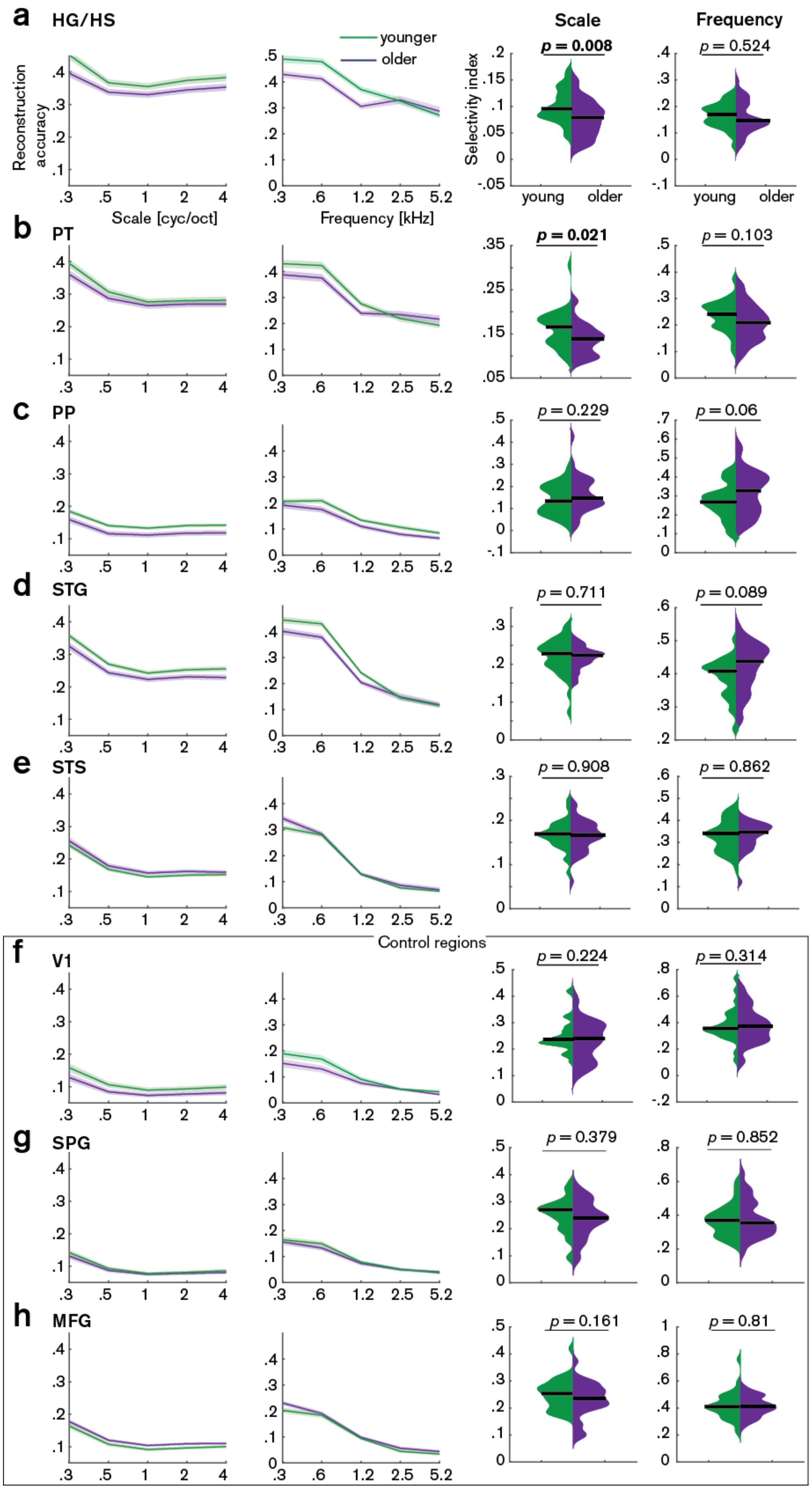

a
HG/HS
Scale
Frequency
younger
older
Reconstruction accuracy
Selectivity index
p = 0.008
p = 0.524
young
older
Scale [cyc/oct]
Frequency [kHz]
b
PT
p = 0.021
p = 0.103
c
PP
p = 0.229
p = 0.06
d
STG
p = 0.711
p = 0.089
e
STS
p = 0.908
p = 0.862
Control regions
f
V1
p = 0.224
p = 0.314
g
SPG
p = 0.379
p = 0.852
h
MFG
p = 0.161
p = 0.81

**Figure 7 – figure supplement 3. Decoding of spectral scale and frequency in ROIs.** Mean (±SE) reconstruction accuracy profiles (left) and selectivity index (right) per age group for spectral scale and frequency in (a) Heschl's gyrus and sulcus (HG/HS), (b) planum temporale (PT), (c) planum polare (PP), (d) superior temporal gyrus (STG), (e) superior temporal sulcus (STS) and the control regions (f) calcarine sulcus (V1), (g) superior parietal gyrus (SPG) and middle frontal gyrus (MFG). Selectivity index was compared per ROI between age groups using exact permutation test. Black line indicates the median *SI*. Note that removing the young outlier participant (> 2 SD away from the mean) for scale in PT reduces the mean age difference to a still significant *SI* difference = 0.015, $p = 0.026$.
